## Supplementary Figures for "Balance of Activity during a Critical Period Tunes a Developing Network"

### Supplementary Information

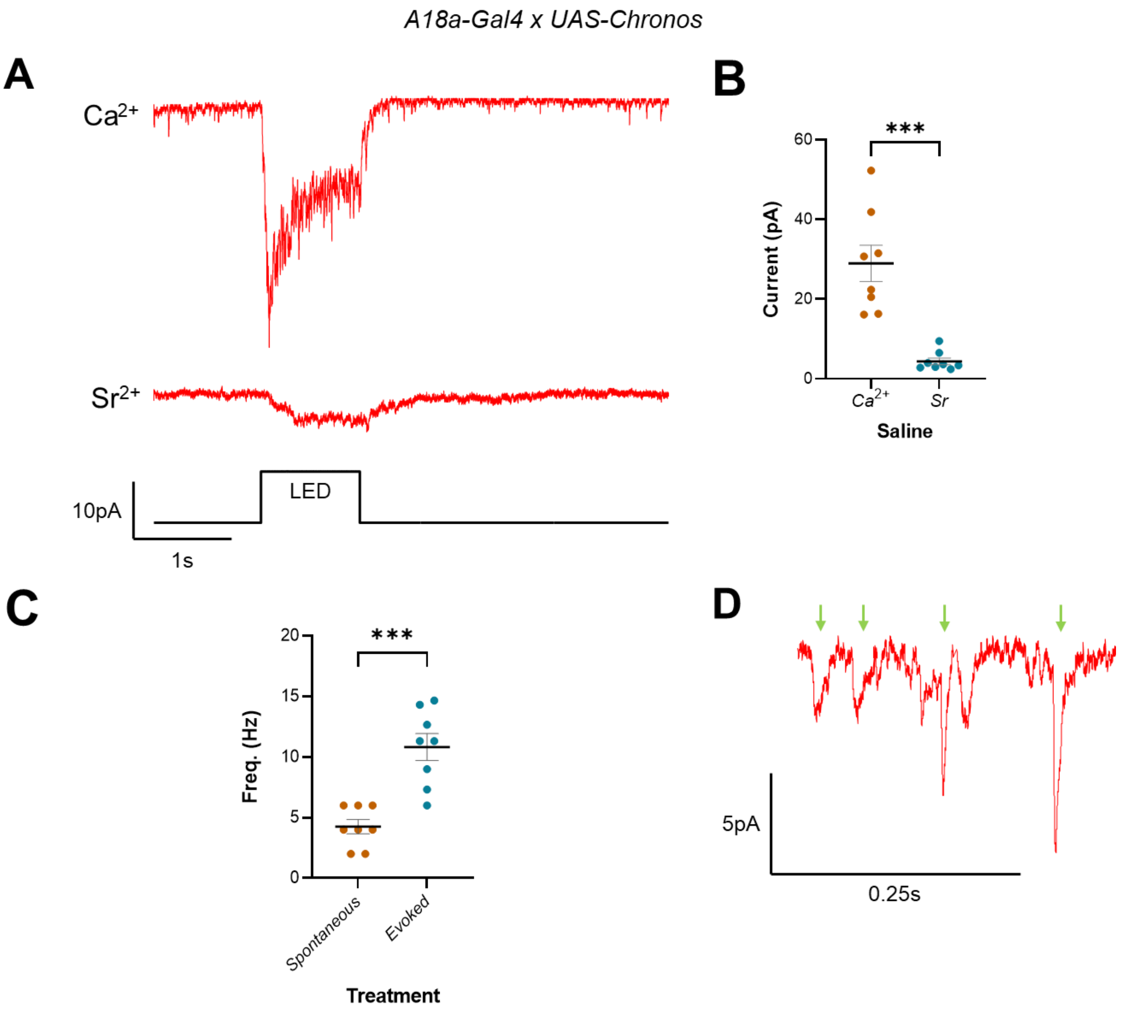


**Figure S1: Optogenetic activation of A18a in 6mM $Sr^{2+}$ saline stimulates asynchronous release of minis.** (A) Representative traces show synaptic currents recorded in aCC following optogenetic stimulation (1s LED) of A18a in control ($Ca^{2+}$) or 6mM $Sr^{2+}$ saline. Recordings made in control saline feature large synaptic events (~30pA), indicative of normal vesicular release from the premotor. By contrast, the large current response to optogenetic stimulation is absent in 6mM $Sr^{2+}$. (B) Analysis of peak currents recorded in aCC, elicited by optogenetic stimulation of A18a are significantly larger in control ($Ca^{2+}$) than 6mM $Sr^{2+}$ saline (28.96 ± 4.56s, $Ca^{2+}$; 4.4 ± 0.85s, $Sr^{2+}$*;* $n$ = 8, 8; $p < 0.0001$), thus, the changes we observed in current amplitude are consistent with the asynchronous mini release reported when using $Sr^{2+}$ saline in mammals (Xu-Friedman and Regehr, 1999; Bekkers and Clements, 1999). (C) Similarly, there is a significant increase in the frequency of asynchronous synaptic events (minis) recorded in aCC in 6mM $Sr^{2+}$ saline, in the 3s following stimulation of A18a (Evoked) versus the 3s prior (4.25 ± 0.59Hz, Spontaneous; 10.83 ± 1.12Hz, Evoked; $n$ = 8, 8; $p < 0.0001$). (D) Minis (green arrows) were defined as ≥2pA events with a short rise time and slow decay. In combination, these results suggest recording current in aCC following optogenetic stimulation of A18a in 6mM $Sr^{2+}$ saline, enables quantifying the frequency and amplitude of minis elicited by this premotor interneuron.

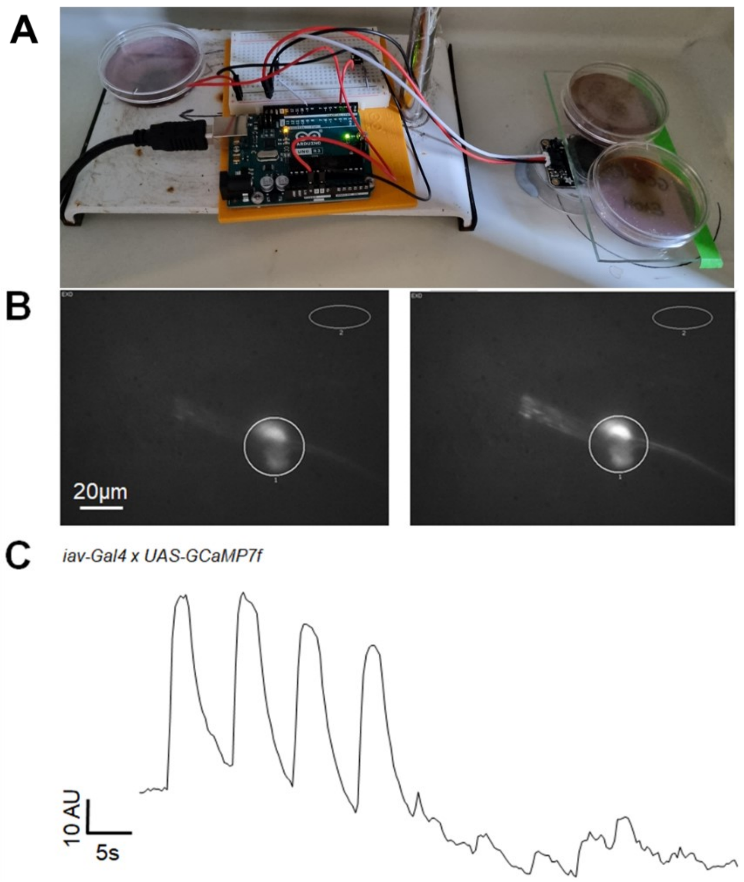


**Figure S2: A custom-built device provides mechanical stimulation of chordotonal neurons.** (A) image of the device, showing an Arduino Uno Rev 3 printed circuit board with microcontroller, that drives a speaker according to timings and frequency (i.e., 17-19h AEL at 0.5Hz) coded using $C^{++}$ in Arduino IDE, and with an amplitude adjusted using a potentiometer. (B) GCaMP imaging demonstrating activity in the chordotonal neurons prior to (left) and following (right) stimulation by the device (response shown is to 1s tone at 80dB). (C) Representative trace of the activity shown in (B).

Bekkers, J. M. and Clements, J. D. (1999) 'Quantal amplitude and quantal variance of strontium-induced asynchronous EPSCs in rat dentate granule neurons', *Journal of Physiology-London,* 516(1), pp. 227-248.

Xu-Friedman, M. A. and Regehr, W. G. (1999) 'Presynaptic strontium dynamics and synaptic transmission', *Biophysical Journal,* 76(4), pp. 2029-2042.
